# Homoploid Hybridization Resolves the Origin of Octoploid Strawberries

**DOI:** 10.1101/2024.09.12.612680

**Authors:** Zhen Fan, Aaron Liston, Douglas Soltis, Pamela Soltis, Tia-Lynn Ashman, Kim Hummer, Vance M. Whitaker

**Author notes:** Corresponding authors: Aaron Liston, Vance M. Whitaker.

## Abstract

The identity of the diploid progenitors of octoploid cultivated strawberry (*Fragaria × ananassa*) has been subject to much debate. Past work identified four subgenomes and consistent evidence for *F. californica* (previously named *F. vesca* subsp. *bracteata*) and *F. iinumae* as donors for subgenomes A and B, respectively, with conflicting results for the origins of subgenomes C and D. Here, reticulate phylogeny and admixture analysis support hybridization between *F. viridis* and *F. vesca* in the ancestry of subgenome A, and between *F. nipponica* and *F. iinumae* in the ancestry of subgenome B. Using an LTR-age-distribution-based approach, we estimate that the octoploid and its intermediate hexaploid and tetraploid ancestors emerged approximately 0.8, 2, and 3 million years ago, respectively. These results provide an explanation for previous reports of *F. viridis* and *F. nipponica* as donors of the C and D subgenomes and unify conflicting hypotheses about the evolutionary origin of octoploid *Fragaria*.

## Main

*Fragaria*, commonly known as strawberry, exhibits a range of ploidy from diploid to decaploid (2n = 2×−10× = 14−70) and occurs across the Northern Hemisphere. The octoploid cultivated strawberry (*Fragaria × ananassa*) is a vitally important fruit crop with steadily increasing consumption ^1^. Although the cultivated strawberry has a short ∼300-year history since its origins via interspecific hybridization between *Fragaria chiloensis* and *Fragaria virginiana*, these two octoploid progenitors evolved through a series of whole-genome duplication events, hybridizations, and subsequent adaptation over millions of years ^2^.

Polyploidy, the condition of possessing more than two complete sets of chromosomes, has been a pivotal mechanism in the diversification and adaptation of many plant species ^3–5^. In strawberries, polyploidy has contributed not only to increased genetic diversity but also to the enhancement of desirable traits such as fruit size, biomass, and resistance to environmental stresses. Unlike some other recent polyploids which experienced recurrent polyploidization ^6^, current phylogenetic evidence supports a single origin of wild octoploid species ^7^. The octoploid strawberry genome has four subgenomes, A, B, C, and D, which originated from four different diploid species. There is scientific consensus that *F. vesca* subsp. *bracteata* (renamed here *F. californica*, see Results and Discussion) and *F. iinumae* served as the donors for subgenomes A and B, respectively, but the origins of subgenomes C and D have been debated ^2,8–10^. Phylogenetic signal from *F. viridis* in the octoploids was first supported by sequences of low-copy genes ^11^. Based on phylogenetic analysis of ortholog sequences from the first chromosome-scale genome assembly of octoploid strawberry, Edger et al.^2^ proposed that *F. viridis* and *F. nipponica* were donors for subgenomes C and D, respectively. Others proposed that subgenomes C and D formed a sister group, sharing an ancestor with *F. iinumae* ^9,12–14^. Based on new evidence from subgenome-specific Kmers, subgenome assignments for the C and D subgenomes have been clarified ^12,14,15^. In this work, we follow the subgenome assignments (Table S1) used in Jin et al. ^12^.

To identify the diploid donors for the two octoploid species (*F. chiloensis* and *F. virginiana*) that are the progenitors of cultivated strawberry, it is essential to survey a wide range of diploid *Fragaria* species and elucidate their phylogenetic relationships. Previous studies using both chloroplast and nuclear genomes supported two major diploid clades, designated as the Southwest China clade (*F. pentaphylla, F. chinensis, F. nubicola, F. daltoniana*, and *F. nilgerrensis*) and the *F. vesca* clade, including *F. vesca* subspecies and *F. mandshurica* ^16–19^. Discordance in the phylogenetic position of *F. iinumae* and *F. viridis* was reported, likely due to past interspecific hybridization and incomplete lineage sorting ^18^. In Europe, hybrids of diploid *F. vesca* and *F. viridis* have been occasionally observed in areas of sympatry ^20^. Likewise, hybrids of *F. iinumae* and *F. nipponica* exist in Japan ^21,22^. Recently, widespread hybridization across diploid *Fragaria* species was revealed; notably, PhyloNet suggested that *F. viridis* may have contributed to the hybrid origination of the lineage of *F. chinensis, F. nipponica, F. nubicola*, and *F. pentaphylla*, and/or the lineage of *F. vesca* and *F. mandshurica* ^18^.

The diploid *Fragaria* that are distributed across North America were previously identified within three subspecies of *F. vesca* ^23^, which are *bracteata, americana*, and *californica*. Phylogenetic analyses of plastomes ^19^ and nuclear microsatellites ^24,25^ demonstrated the distinctiveness of the North American subspecies from Eurasian *F. vesca*, but American diploid *Fragaria* has been underrepresented in phylogenetic studies of the nuclear genome. While the northwestern North American *F. vesca* subsp. *bracteata* contributed the plastid genome ^19^ to the octoploid strawberry, whether plants from the same geographic region also contributed subgenome A remains untested, due to limited sampling in previous studies. Generating whole genome sequences of American diploids will not only help resolve their relationship with *F. vesca*, but also verify their contribution to the octoploid genome. Additionally, a reticulate phylogeny approach can help us understand how homoploid hybridization among diploid species has contributed to *Fragaria* polyploids.

In this study, we have sought to resolve the cryptic signals from *F. viridis* and *F. nipponica* in the octoploid strawberry genome and unify conflicting hypotheses about the evolutionary path to octoploid strawberry. Additionally, using octoploid strawberry as a model, we refined a framework to date the approximate timing of polyploidization events based on the age of insertion of long terminal repeats (LTR).

Using whole genome sequence data, three bifurcating phylogenetic approaches (ASTRAL, ML, and SVDQuartets) provided similar results regarding the origins of the four subgenomes for both octoploid strawberry species (Fig 1A, Fig. S1 and S2). These analyses showed that subgenome A is sister to a clade of North American diploids including both *F. vesca* subsp. *bracteata* and subsp. *americana*, subgenome B is sister to *F. iinumae*, and subgenomes C and D form a clade with subgenome B and *F. iinumae*, confirming multiple reports ^8,9,12^. In all three trees, two *F. viridis* samples, *F. nilgerrensis*, and *F. nipponica* consistently formed a monophyletic group, sister to the clade of subgenomes B, C, D, and *F. iinumae* clade. Based on their phylogenetic sister relationship with *F. mandshurica* determined by both genomic and plastid DNA sequence data ^7^, along with their distinct morphological characteristics and geographic distributions, the North American diploids (including *F. vesca* subsp. *bracteata*, subsp. *americana*, and subsp. *californica*) are here designated as a single species *Fragaria californica* Cham. & Schltdl. (1827), the oldest available name. Whether infraspecific taxa are warranted will require more intensive sampling across the broad geographic range of the newly recognized *F. californica*. While the plastid genome ^19^ indicates a northwest North American maternal ancestry of the octoploid strawberry, our nuclear phylogenomic results do not resolve where in North America subgenome A originated.

**Figure 1.**
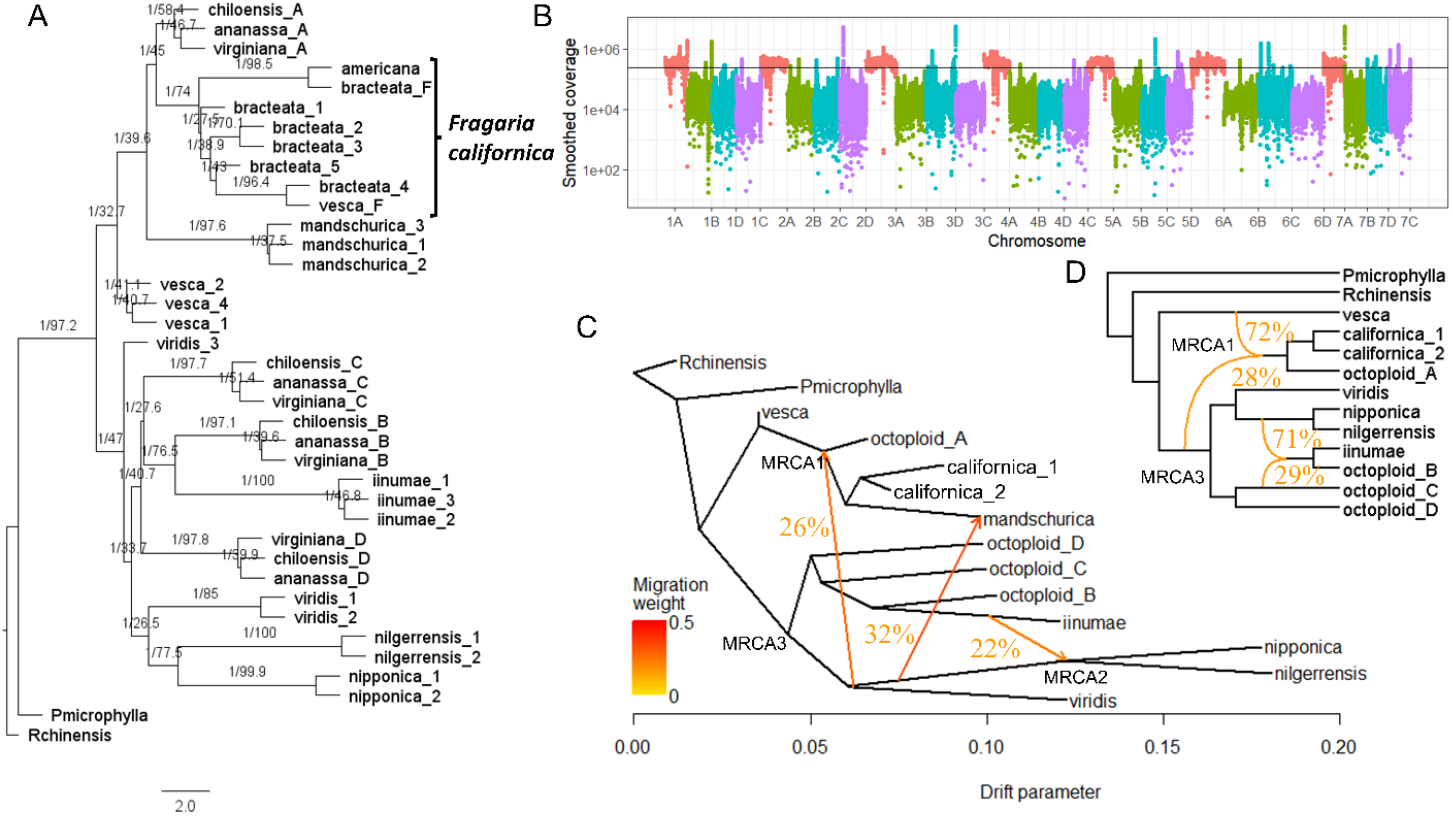
Evidence of homoploid hybridization in the formation of the ancestral octoploid strawberry genome. (A) ASTRAL species tree inferred from 1611 10K-variant-window trees. Branches are annotated by local posterior probabilities and gene concordance factors (%). Pmicrophylla and Rchinensis represent *Potentilla microphylla* and *Rosa chinensis*, which are both outgroups. Four subgenomes in the octoploid species are postfixed with A/B/C/D. (B) Mapping coverage of reads from *F. californica* (*F. vesca* subsp. *bracteata)* against the octoploid genome. The horizonal line indicates the 20% quantile of coverage across subgenome A. (C) TreeMix phylogenetic network. Directional migration edges are annotated by their migration weight (inheritance percentage). (D) Phylogenetic network inferred by Maximum Pseudo-likelihood method. *F. mandshurica* samples were removed due to its recent hybridization to reduce network complexity.

In polyploids, homoeologous exchanges (HEs) and homoeologous recombination (HR) can reshuffle the genetic differences among subgenomes, leading to the admixture of ancestral information within a subgenome. Edger et al.^2^ suggested that extensive HEs from subgenome A contributed 0.5 to 28.3% of the chromosomes in the other subgenomes. However, no signals for large-scale segmental HE were revealed by subsequent Kmer analysis ^14^. Given the proposed biased pattern of HEs from subgenome A to the other subgenomes, our analysis focused on detecting these specific signals. Although three four-taxon trees representing the relationships among the four subgenomes and *F. californica* (Fig. S3) did not reveal significant deviations from expected relationships based on genome-wide SNPs, positive f-branch values between subgenomes B and A (fbranch = 1.67%) and between subgenomes C and A (1.08%) suggested the presence of HE (Fig. S4). To identify regions that underwent HE, we used an alignment-based approach with newly generated whole genome sequences of *F. californica*. We identified 42 candidate HE regions in the B, C, and D subgenomes, totaling 2.7 Mb (≈ 0.34% of the genome, ≈ 0.46% of the B, C, and D subgenomes), which showed high mapping coverage of *F. californica* sequences (Fig. 1B). The higher percentage of overlap with intact TEs (10.7%) compared to the genome-wide average (7.7%) (Fig. S5) suggests the potential inclusion of transposon-derived duplicates, despite applying a minimum size selection of 20 kb. To confirm HE in these regions, ML trees constructed using concatenated SNPs were analyzed. A sister-group relationship of subgenome A or *F. californica* instead of *F. iinumae* with one of the subgenomes B, C, or D was found in 32 regions, supporting their HE assignment (Table S2). In four cases, including the largest identified HE (farr1_chr_2D/2C:3820000-4200000bp), disparities in tree topologies between the two wild octoploid species (*F. chiloensis* and *F. virginiana*) suggest that these HE occurred after the divergence of the two octoploid species, distinguishing them from hybridization among diploids prior to octoploidization. Lastly, for the largest HE, differences in synteny between *F. chiloensis* and *F. virginiana* vs. *F. vesca* corroborate occurrence of this HE after divergence of the two octoploid species (Fig. S6). Our results validated HE occurrence during evolution of octoploid strawberry, though its scale was much smaller than previous estimates ^2,26^. This is likely a result of using reads from the direct donor species of the A subgenome and the exclusion of transposon-derived duplications.

Although *F. californica* and *F. iinumae* were the only species recognized as extant diploid progenitors of the ancestral octoploid strawberry in this and multiple previous studies, elevated mapping coverage of *F. viridis* and *F. nipponica* sequences to the octoploid genome ^12,13^ and a sister relationship with octoploid orthologs in a substantial number of gene trees were often observed ^2,10,11^. Therefore, multiple phylogenetic network approaches (Treemix, admixture graph and Phylonet) were used to infer hybridization during the evolution of the octoploid species. Three hybridization events were determined as the optimal number in Treemix ^27^. The inferred hybridization edges were from *F. viridis* to the MRCA (MRCA1) of subgenome A of the octoploid species, *F. californica*, and *F. mandschurica* (Migration weight = 26.1%); from the MRCA (MRCA2) of *F. nipponica* and *F. nilgerrensis* to *F. mandschurica* (32.73%); and from *F. iinumae* to MRCA2 (21.6%) (Fig. 1C). The admixture graph ^28^ estimated that introgression from *F. viridis* contributed 24% (CI: 13%-37%) to the genome of MRCA1. The other two admixture proportions were 26% (CI: 10%-38%, MRCA2 to *F. mandschurica*) and 28% (CI: 11%-96%, *F. iinumae* to MRCA2), respectively (Fig. S7). PhyloNet results corroborated that MRCA1 was a hybrid, but its second parental species was assigned to MRCA3 (Fig. 1D, 28%), which is the MRCA of all sampled diploid species except the *F. vesca* clade. In contrast, the Phylonet analysis suggests that the ancestor of *F. iinumae* and subgenome B could be a hybrid or that *F. iinumae* has introgressed into both *F. nipponica* and the diploid donor of subgenome C.

Therefore, the phylogenetic sister relationship of octoploid and *F. viridis* orthologs and higher mapping rates of *F. viridis* to the octoploid genomes appear to result from its introgression into the ancestor of subgenome A prior to octoploidization. On the other hand, the close relationship of *F. nipponica* and octoploids in gene trees is likely due to its hybridization with *F. iinumae*. These results reconcile the conflicting phylogenetic hypotheses for the origins of the octoploid subgenomes ^2,9,10^.

Both Ks-based and phylogeny-based age estimation ^29^ are unsuitable for octoploid strawberry due to its recent genome duplication and the presumed extinction of the C and D diploid donors ^8,30^. Recently, LTR age estimation has been used to provide a range of values for the age of polyploids ^31^. LTR movement occurs continuously, acting like timestamp embedded in the genome, and specific LTRs are active at different times and in even closely related species ^32,33^. Therefore, the timing of LTR insertions that uniquely distributed in individual subgenomes provided a way to date polyploidization events. For octoploid strawberry, Session & Rokhsar ^14^ estimated that hexaploidization occurred around 3 million years ago (MYA) based on the peak in age distribution for common LTRs shared by the B, C, and D subgenomes. Our approach seeks to refine these estimates by providing estimates and confidence intervals (CI) for the ages of the octoploid, intermediate hexaploid and tetraploid based on LTR age, as explained below.

All subgenome donors (A, B, C, D) of octoploid strawberry share a common ancestor. Once they diverged, each diploid species began accumulating species-specific LTRs in their genomes until they hybridized into a polyploid species. The polyploidizations that formed octoploid strawberry occurred in a specific sequence: tetraploid (C and D subgenomes), hexaploid (B, C and D subgenomes), and then octoploid (Fig. 2A) ^14^. The timing of the hybridization event between the C and D donors can be dated using the insertion time of the youngest C- and D-specific LTRs (Fig. 2B). After this hybridization event, in the tetraploid ancestor, LTR movement was random, allowing the same LTR to insert into either the C or D subgenome. As a result, the C and D subgenomes share LTRs that were active in the tetraploid ancestor; the youngest LTRs in the intermediate tetraploid and subgenome B donor can be used to determine the age of hexaploid formation (Fig. 2B). The same process applies when the tetraploid ancestor hybridized with the B subgenome donor, at which point newer LTRs began to be shared among the B, C, and D subgenomes. Using 5% quantiles of these LTR age distributions and six independent genome assemblies of octoploid species, we could determine the age and its confidence interval (CI) for each polyploidization event. The LTR age distributions confirmed the order of polyploidization, with the LTRs specific to subgenomes C and D showing the oldest distribution, whereas subgenome A and the intermediate hexaploid showed the youngest (Fig. 2C, Supplementary Data1). The LTR-based approach estimated that tetraploidization between the C and D subgenome donors occurred approximately 3.0 MYA (CI = [4.5, 1.2]), hexaploidization with the B subgenome donor at roughly 2.2 MYA (CI = [2.7, 1.8]), and octoploidization with the A subgenome donor at 0.8 MYA (CI = [1.0, 0.4]) (Fig. 2D). Because the LTR approach is not influenced by the absence of diploid donors and hybridization prior to polyploidization, its estimates were lower than those from phylogeny-based approaches (Table 1). Our age estimation for the formation of the octoploid is close to the estimate based on a calibrated tree inferred from plastid DNA sequences (1MYA) ^7^. The CIs for both hexaploid and octoploid formation were less than 1 MYA. However, for the most ancient polyploidization, the retention of intact subgenome-specific TEs was low, especially for subgenome C, resulting in a wide CI for tetraploid formation.

**Table 1.** Phylogenetic dating of octoploid and intermediate polyploid ancestors of octoploid strawberry.

| Formation | LTR age-based approach | Phylogeny-based approach |  |  |  |  |  |
| --- | --- | --- | --- | --- | --- | --- | --- |
|  |  | Filtered single-copy orthologs<sup>a</sup> |  |  | All single-copy orthologs |  |  |
|  |  | R8s<sup>b</sup> | LSD2 | RelTime | R8s | LSD2 | RelTime |
| Octoploid | 0.8<br>(1.0, 0.4)<sup>c</sup> | 2.74<br>(2.80, 2.67) | 1.91<br>(2.61, 1.51) | 1.44 | 3.33<br>(3.42, 3.22) | 2.23<br>(2.91, 1.85) | 1.73 |
| Hexaploid | 2.2<br>(2.7, 1.8) | 6.74<br>(6.87, 6.60) | 5.10<br>(6.33, 4.02) | 3.30 | 6.27<br>(6.39, 6.18) | 5.29<br>(6.93, 4.20) | 3.41 |
| Tetraploid | 3.0<br>(4.5, 1.2) | 7.91<br>(8.05, 7.75) | 6.24<br>(7.97, 5.18) | 4.06 | 7.18<br>(7.27, 7.08) | 6.32<br>(8.23, 5.19) | 4.10 |

<sup>a</sup> Orthologs which showed potential admixture signals based on four-taxon trees were removed.
<sup>b</sup> R8s, LSD2, and RelTime are three popular methods for dating evolutionary divergences.
<sup>c</sup> Confidence intervals are given.

**Figure 2.**
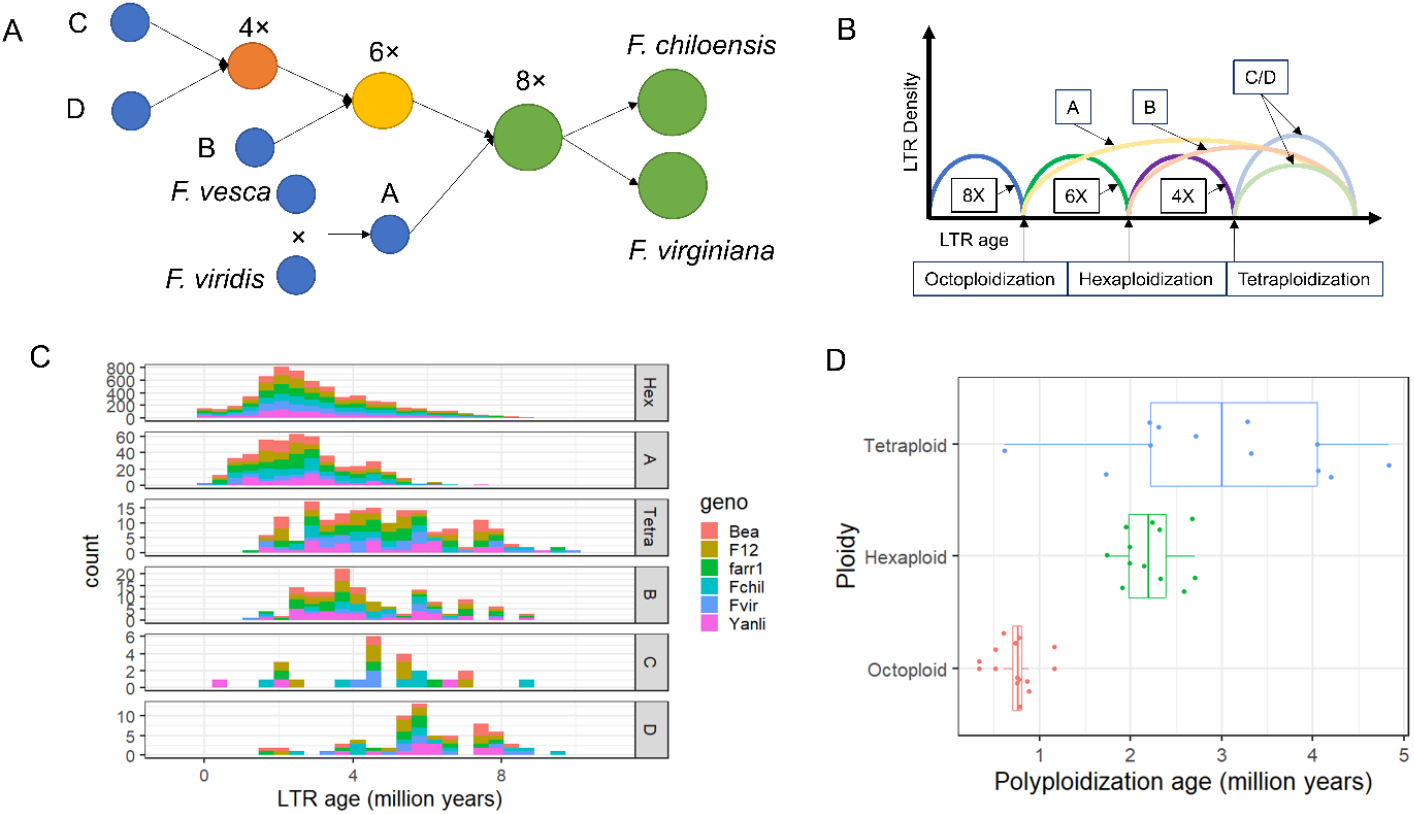
Dating the approximate timing of polyploidization in the formation of octoploid strawberry. (A) Schematics of evolutionary path to octoploid *Fragaria*. (B) Theoretical LTR age distributions in subgenome donors (A, B, C, and D) and intermediate polyploid ancestors (Intermediate tetraploid: 4X, Intermediate tetraploid: 6X, octoploid: 8X). (C) Cumulative LTR age distributions in diploid subgenome donors (A, B, C and D) and intermediate hexaploid (Hex) and tetraploid (Tetra) ancestors across six genomes. Bea, F12, farr1 and Yanili are genome assemblies of *F. × ananassa* accessions. Fchil and Fvir are genomes of *F. chiloensis* and *F. virginiana*, respectively. (D) Boxplots of age estimates for polyploidization events. Each dot represents one estimate based on one LTR distribution of one genome assembly. A total of 12 data points is used to date each polyploidization event.

These age estimates are consistent with the earliest fossil records of *Fragaria* from the Late Pliocene (3.6-2.6 Ma) of the Canadian Arctic and Yunnan, China. The Canadian record was based on a single well-preserved achene, but no image or specimen is available ^20,34^. The Chinese record is based on 19 well-preserved achenes that are vouchered and photographed ^35^. A Miocene report of *Fragaria* is considered unreliable, as it compares the fruiting structure to both *Fragaria vesca* and *Potentilla indica* (as *Fragaria indica*) ^36^.

It has been hypothesized that these sequential allopolyploidization events 3.0-0.8 MYA leading to the origin of octoploid *Fragaria* occurred in Beringia ^30^. During this time frame, Beringia is hypothesized to have been characterized by mixed conifer forests with a diverse herbaceous understory ^37^, appropriate habitat for *Fragaria*. Whether the homoploid hybridization events inferred here also occurred in Beringia is unknown. Environmental DNA (eDNA) has documented over 100 different plant genera from a 2-million-year-old ecosystem in northern Greenland ^38^; eDNA studies in the former Beringia could potentially provide physical evidence bearing on the evolutionary history proposed here.

In summary, based on admixture analyses and reticulate phylogeny, we propose that homoploid hybridization between *F. viridis* and *F. vesca* led to the formation of the North American *F. californica* and subgenome A of octoploid strawberry. Hybridization among *F. iinumae, F. nipponica*, and possibly the donor of subgenome C might have resulted in the sister relationship of *F. nipponica* and octoploid species in gene trees. Homoeologous exchange contributed to genome reshuffling but only on a small scale. These findings resolve conflicting hypotheses about the evolutionary origin of octoploid strawberry and will guide efforts to enhance genetic diversity in *F. × ananassa* from diploid species through interspecific hybrid and horizontal gene transfer. Our framework for dating polyploidization events using the 5% quantiles of LTR age distributions in intermediate polyploids and diploid donors produced estimated ages of octoploid, intermediate hexaploid, and tetraploid formation in the octoploid strawberry as 0.8, 2, and 3 MYA, respectively. This approach, which does not require sequences from extant ancestral diploid species, provides a valuable addition to existing methods for dating polyploidization events when subgenome donor species are unavailable.

## Materials and methods

### Sample collection and sequencing

Freeze-dried leaves of ten diploid *Fragaria* samples, including five *F. vesca* subsp. *bracteata* and one *F. vesca* subsp. *americana* samples (renamed as *F. californica* based on results), were used for CTAB DNA extraction. The geographic location and taxon designation of each sample are provided in a supplementary table (Table S3), and voucher specimens are deposited in the Oregon State University Herbarium (OSC). DNA was sent to Novogene Corporation Inc., Sacramento, CA, USA, for whole genome sequencing (WGS). Paired-end (2×150 bp) libraries were constructed and sequenced on Illumina HiSeq X Ten platform. Short reads of 16 additional diploid *Fragaria* samples and two outgroup species (*Potentilla microphylla* and *Rosa chinensis*) were obtained from six previous studies ^2,17,18,39–41^. Assembles for each subgenome in *F. × ananassa* ^42^, *F. chiloensis* and *F. virginiana* ^12^ were extracted from the whole genome assembles based on the most recent subgenome assignment ^12^. Simulated 2×150 bp pair-end short reads for them were generated using ART V20160605 ^43^. Variant calling follows our previous work ^44^. Briefly, reads were aligned to a chromosome-scale *F. vesca* genome assembly V6.0 ^45^ using SNAP V2.0.3. SNPs and indels were called and filtered using GATK V4.4.0 ^46^.

### Species trees

A total of 16,114,631 SNPs and 40 terminal nodes (Table S3) including all *Fragaria* species related to origins of the octoploid species were used to reconstruct the phylogeny of *Fragaria*. Three approaches were applied: (1) SNPs were grouped into 1611 windows with 10,000 variants in each. A maximum likelihood (ML) tree was constructed for each window using IQ-TREE V2.3 ^47^. ASTRAL V5.15 ^48^ was used with default settings to construct a species tree with these 1611 trees as input; (2) an SVDQuartets ^49^ tree was built using the same 1611 partitions with paup V4.0; (3) a concatenated tree was inferred with all SNPs using IQ-TREE. Two previously identified *F. californica* samples (bracteata_E1 and bracteata_E2) were found to be octoploid samples based on uniform read-coverage distributions across four subgenomes and were thus removed from the following analyses.

### Admixture analyses and homoeologous exchange identification

To investigate percentages of genomic admixture, Fbranch values were obtained using Dsuite V0.4 ^50^. To identify genomic regions of HE, WGS of five *F. californica* samples were aligned to the octoploid genome ^42^. Read coverage for every 10-kb window was obtained using the bedcov function in samtools V1.19 ^51^. Median values across samples were smoothed using the SS function (m=1, spar = 0.05) in R npreg library ^52^. The 20% quantile of read coverage within subgenome A was used as the cutoff to identify HE regions in other subgenomes. Adjacent regions with a gap of less than 30 kb were merged. Potential HE regions smaller than 20 kb were pruned to remove signals of recent transposon insertions. Syntenic regions in the *F. vesca* genome V6.0 were inferred using CoGe SynMap (https://genomevolution.org/coge/). Syntenic regions were identified for 39 of 42 regions in the *F. vesca* genome ^45^. A ML tree was inferred for each individual HE region. The topology of each tree was manually examined to validate HE.

### Reticulate networks

The first reticulate tree was built using TreeMix V1.12 ^53^ for 10 species and four subgenomes of *F. virginiana* and *F. chiloensis*. Two or three samples of each taxon were used (Table S4). The optimal number of reticulate edges was determined using OptM ^27^. An admixture graph ^28^ was built using Admixtools2, starting with the best TreeMix tree topology. Automated searches for better topologies with higher likelihoods did not yield improved networks. Genomic admixture percentages for the migration edges were calculated using f2 values. The third species network was constructed using Phylonet ^54^, with the same samples and taxa as the TreeMix input, except that the *F. mandshurica* samples were removed to reduce model complexity. Maximum pseudo-likelihood networks were built with one to four reticulation nodes over 100 rounds of searching using 1611 10K-variant window trees. The network with two reticulate nodes showed a 5% improvement in likelihood over the network with one node, while the improvement dropped to 1% for three reticulate nodes (Table S5). Thus, two reticulate nodes were chosen as the best fit, consistent with the TreeMix results after the exclusion of the recent hybrid *F. mandshurica*. In a different run, a different network achieved a similar likelihood (log likelihood = -2022481) as our best network but identified MRCA3 as the hybrid of MRCA1 (Fig. 1). Given the large diversity of diploid *Fragaria* species in Asia, the older estimated age of MRCA3 compared to MRCA1 (Qiao et al., 2021), and the challenges in determining the direction of hybridization ^55^, the network shown in Figure 1D appears more plausible.

### Dating of polyploidization events

High-quality genome assemblies of one *F. chiloensis*, one *F. virginiana* ^12^, and four *F. × ananassa* haplotypes ^42,56,57^ were downloaded from GDR (https://www.rosaceae.org/). Each assembly was separated into four subgenome-specific assemblies. Repetitive 15-mers were counted using KMC V3.2 ^58^ and then filtered with minimum frequency of 100. To obtain repetitive 15-mers specific to the intermediate tetraploid and hexaploid *Fragaria*, repetitive 15-mer sets for subgenomes C and D and subgenomes B, C, and D were first intersected, respectively. Then all 15-mers of subgenome A and B, and subgenome A were removed from tetraploid and hexaploid repetitive 15-mer sets, respectively. The diploid donor-specific 15-mers were obtained by removing all 15-mers of other subgenomes from each of the subgenomic repetitive 15-mers sets. The 15-mer sets were aligned to their own octoploid genome using bowtie ^59^ with parameters (-S -v 0). Intact TEs and their insertion time for each haplotype assembly were identified and computed using EDTA V2.1 ^60^. The rate of nucleotide substitution was set to 0.7 × 10^-8 sub/year according to previous calibration adjusted for the time of *Fragaria* MRCA at 8 MYA ^14^. Intact LTRs overlapping with each of the 15-mer sets were extracted. A minimum coverage of 1% of the LTR and two overlapping 15-mers were used to filter the overlapping LTRs. 5% quantiles were obtained for each 15-mer set. To date polyploids based on phylogenetic trees, an ASTRAL tree was built based on 1307 single-copy orthologs. The orthologs were identified in ten diploid assemblies and four subgenomes in both *F. chiloensis* and *F. virginiana* using OrthoFinder ^61^. MAFFT ^62^ and IQ-Tree were used to align orthologs and infer gene trees. Three popular phylogenetic dating methods, R8s ^63^ , LSD2 ^64^ and RelTime ^65^ were applied. Two fossils were used for calibration: The *Rosa* fossil calibration is based on an Early Eocene (55.8 - 48.6 Ma) mold/impression fossil from Idaho, U.S.A. ^66^, confirmed by Bruce Tiffney (paleobiodb.org). The *Potentilla* fossil calibration is based on an Oligocene (33.9-23.0 Ma) mold/impression fossil from Montana, U.S.A. ^67^ confirmed by Hallie Sims (paleobiodb.org). PL and TN method with 100 bootstrap datasets was applied in R8s to obtain confidence interval for divergence time. *Potentilla microphylla* (23 MYA) fossil was applied as fixed age in R8s. The substitution model “JTT+F+I+R8” was set in LSD2. To reduce the effect of *F. viridis* introgression into *F. vesca* and the absence of *F. californica* genome, dating was also inferred using non-admixed gene trees filtered by a four-taxon tree (*P. microphylla, F. viridis, (F. vesca, F. virginiana* subgenome A)). ASTRAL was used to build a tree from this non-admixed gene set as the input for dating.

## Data Availability

All codes are deposited in Github (https://github.com/zhen0506/Strawberry-Homoploid-Hybridization-). Supplementary data include species trees, raw input and output of phylogenetic dating software and SNP database is available in Zenodo (10.5281/zenodo.13513299). Raw sequencing data is available in NCBI (PRJNA1153529).

